# Endolysosomal degradation of Tau and its role in glucocorticoid-driven hippocampal malfunction

**DOI:** 10.1101/352724

**Authors:** João Vaz-Silva, Mei Zhu, Qi Jin, Viktoriya Zhuravleva, Patrícia Gomes, Sebastian Quintremil, Torcato Meira, Joana Silva, Chrysoula Dioli, Carina Soares-Cunha, Nikolaos P. Daskalakis, Nuno Sousa, Ioannis Sotiropoulos, Clarissa L Waites

## Abstract

Emerging studies implicate Tau as an essential mediator of neuronal atrophy and cognitive impairment in Alzheimer’s disease (AD), yet the factors that precipitate Tau dysfunction in AD are poorly understood. Chronic environmental stress and elevated glucocorticoids (GC), the major stress hormones, are associated with increased risk of AD, and have been shown to trigger intracellular Tau accumulation and downstream Tau-dependent neuronal dysfunction. However, the mechanisms through which stress and GC disrupt Tau clearance and degradation in neurons remain unclear. Here, we demonstrate that Tau undergoes degradation via endolysosomal sorting in a pathway requiring the small GTPase Rab35 and the endosomal sorting complex required for transport (ESCRT) machinery. Furthermore, we find that GC impair Tau degradation by decreasing Rab35 levels, and that AAV-mediated expression of Rab35 in the hippocampus rescues GC-induced Tau accumulation and related neurostructural deficits. These studies indicate that the Rab35/ESCRT pathway is essential for Tau clearance and part of the mechanism through which GC precipitate brain pathology.

## Introduction

Intraneuronal accumulation of Tau protein represents a central pathogenic process in Alzheimer’s disease (AD), and is hypothesized to mediate the detrimental effects of amyloid-beta (Aβ) on neuronal function and cognitive performance (Guo, Noble et al., 2017, Ittner, Ke et al., 2010, Roberson, Scearce-Levie et al., 2007). Tau accumulation and hyperphosphorylation is linked to synaptic atrophy, neuronal dysfunction, and cognitive deficits (Kimura, Yamashita et al., 2007, Yin, Gao et al., 2016), and triggers these events by disrupting axonal trafficking (Vossel, Zhang et al., 2010), promoting GluN2B-related excitotoxicity (Ittner et al., 2010, Zempel, Thies et al., 2010), and suppressing nuclear CREB-mediated synthesis of synapse- and memory-related proteins (Yin et al., 2016), among other effects. Furthermore, emerging studies support a crucial role for Tau in diverse brain pathologies (for review, see (Sotiropoulos, Galas et al., 2017)) including prolonged exposure to stressful conditions, a known risk factor for AD and major depressive disorder (Vyas, Rodrigues et al., 2016). In particular, recent studies demonstrate that exposure to chronic environmental stress or the major stress hormones, glucocorticoids (GC), triggers the accumulation of Tau and its synaptic missorting, precipitating dendritic atrophy and synaptic dysfunction in a Tau-dependent manner (Dioli, Patricio et al., 2017, Lopes, Lopes et al., 2016a, Lopes, Teplytska et al., 2016b, Pallas-Bazarra, Jurado-Arjona et al., 2016, Pinheiro, Silva et al., 2015). However, the cellular mechanisms responsible for stress/GC-induced accumulation of Tau remain unclear.

A major cause of Tau accumulation is the dysfunction of its degradative pathways (Khanna, Kovalevich et al., 2016). Tau degradation has been shown to occur through two distinct mechanisms, the ubiquitin-proteasome system and the autophagy-lysosome pathway (Lee, Lee et al., 2013, Zhang, Chen et al., 2017). Indeed, proteasome and lysosome inhibitors can delay Tau turnover and promote Tau-driven neuropathology (Hamano, Gendron et al., 2008, Zhang, Liu et al., 2005). However, dysfunction of a third degradative pathway, the endolysosomal system, is also linked to AD and other neurodegenerative conditions that exhibit Tau accumulation, including Parkinson’s disease (Kett & Dauer, 2016, Rivero-Rios, Gomez-Suaga et al., 2015, Small, 2017). Nevertheless, the role of the endolysosomal pathway in the clearance of Tau is almost completely unexplored.

Here, we report that the small GTPase Rab35 and the endosomal sorting complex required for transport (ESCRT) machinery mediate the delivery of Tau to lysosomes via early endosomes and multivesicular bodies (MVBs). We further demonstrate that this pathway is negatively regulated by GC, which suppress Rab35 transcription, thereby inhibiting Tau degradation and promoting its accumulation. Finally, we show that *in vitro* and *in vivo* overexpression of Rab35 can rescue GC-induced Tau accumulation and neurostructural deficits in hippocampal neurons. These findings demonstrate Rab35’s critical role as a regulator of Tau clearance under physiological and pathological conditions.

## Results

### The ESCRT pathway is necessary and sufficient for Tau degradation

Despite its primarily axonal and microtubule-related distribution, Tau is also found in the somatodendritic compartment and associated with lipid membranes (Georgieva, Xiao et al., 2014, Pooler & Hanger, 2010). To determine whether Tau undergoes endosomal sorting, we looked for Tau immunoreactivity in multivesicular bodies (MVBs), late endosomal structures that deliver cargo to lysosomes. Immuno-electron microscopy in rat hippocampal tissue revealed MVBs that contain Tau immunogold labeling (Fig. 1A; Fig. EV1A), indicating that Tau is sorted into MVBs and can potentially undergo degradation via the endocytic pathway. MVB biogenesis is catalyzed by the endosomal sorting complex required for transport (ESCRT) system, a series of protein complexes that recruit ubiquitylated cargo destined for degradation and facilitate the formation of intralumenal vesicles (Raiborg & Stenmark, 2009). The initial component of the ESCRT-0 complex, Hrs, is essential for binding and recruitment of ubiquitylated cargo into MVBs (Frankel & Audhya, 2017, Raiborg & Stenmark, 2009). To examine whether Hrs interacts with Tau, we performed the proximity ligation assay (PLA; see Fig. 1B) in N2a cells overexpressing mCh-tagged Hrs (Fig. 1C; Fig. EV1B). Here, we detected fluorescent puncta, representing close (20–50 nm) proximity of Hrs and Tau, that were associated with Hrs-positive early endosomes (Fig. 1C) and increased upon overexpression of GFP-tagged wild-type Tau (GFP-wtTau; 4R0N isoform; Hoover, Reed et al., 2010)(Fig. EV1C-D), demonstrating specificity of the PLA signal for the Hrs/Tau interaction. This interaction was confirmed by co-immunoprecipitation (IP) experiments in N2a cells overexpressing FLAG-Hrs and either GFP or GFP-wtTau, wherein Hrs was specifically pulled down by GFP-wtTau (Fig. EV1E). Treatment with a deubiquitylating enzyme (DUB) inhibitor to increase ubiquitylated Tau species further increased the Hrs/Tau interaction as measured by both co-IP and PLA (Fig. EV1F-I), indicating that ubiquitylation promotes Tau’s association with Hrs and subsequent entry into ESCRT pathway. Using super-resolution fluorescence microscopy, we also identified Tau in both membranes and lumen of Hrs-, EEA1-, and Rab5-positive early endosomes in N2a cells (Fig. 1D-E). These findings demonstrate the presence of Tau in endosomal compartments at both early and late stages of the endolysosomal pathway.

**Figure 1.**
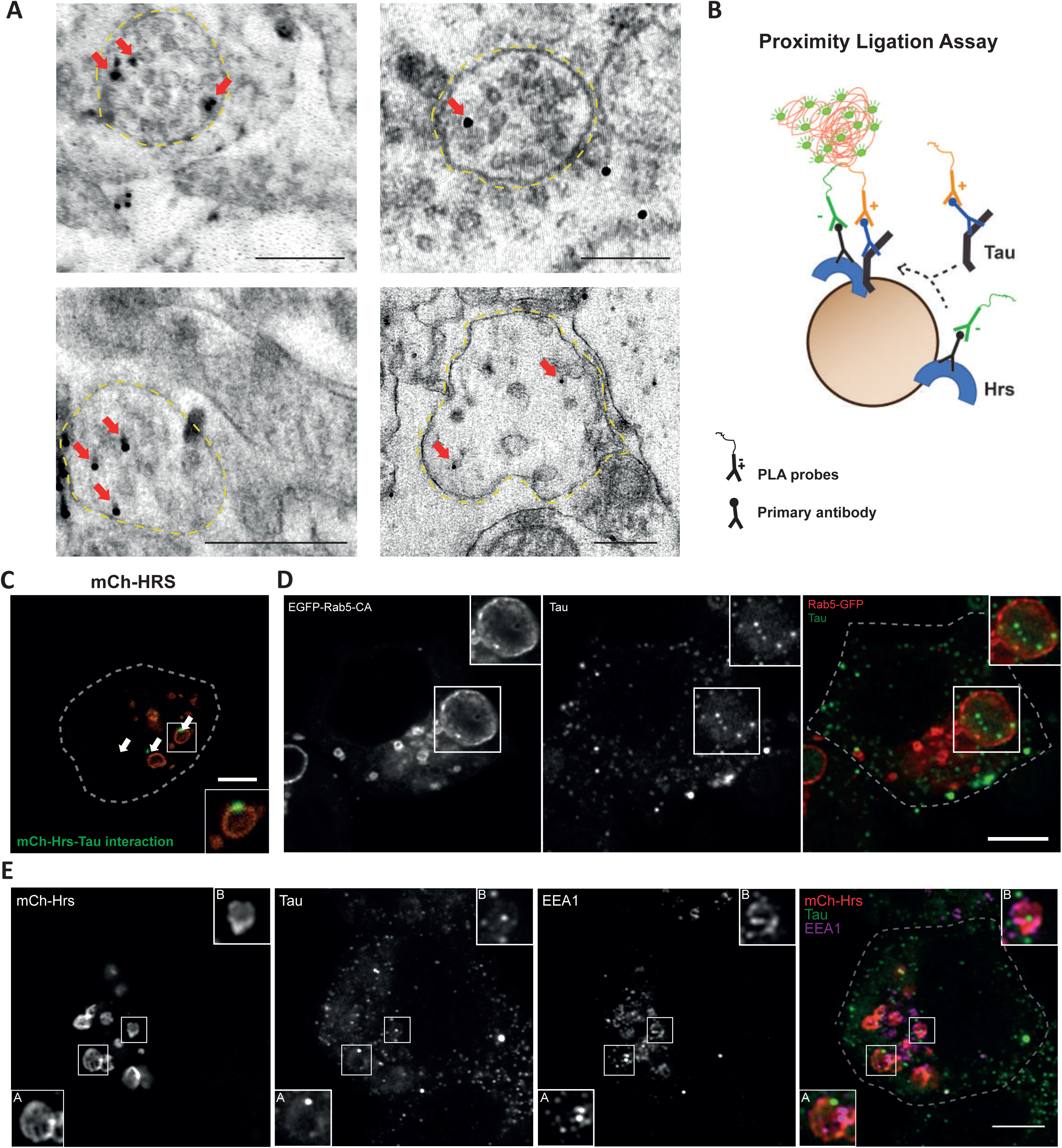
Tau protein localizes to early endosomes and MVBs. (A) Electron micrographs of immunogold-labeled Tau (arrows) in multivesicular bodies (yellow dashed lines) of rat hippocampus (scale bar: 250nm). (B) Schematic diagram of proximity ligation assay (PLA), wherein primary antibodies against Hrs or Tau are detected by secondary antibodies conjugated to DNA. Ligation and amplification of DNA, resulting in green signal, only occurs when proteins are in close (20–50 nm) proximity. (C) PLA signal (green) for Hrs/Tau interaction (arrows) on an mCh-Hrs-labeled endosome (red) in N2a cells; scale bar: 5 μm (see also Fig. S1). (D) Super-resolution images of N2a cells transfected with constitutively-active Rab5 (EGFP-Rab5-CA) and immunostained for Tau, revealing the presence of Tau in both membranes and lumen of early endosomes (scale bar: 5 μm). (E) Super-resolution images of N2a cells transfected with mCh-Hrs and immunostained for Tau and EEA1. Higher-magnification insets (A and B) show Hrs- and EEA1-positive endosomes with juxtaposed Tau puncta present both in the membrane and lumen; scale bar: 5 μm.

### Rab35 promotes Tau degradation through the ESCRT pathway

To assess the role of the ESCRT pathway in Tau clearance (Fig. 2A), we measured Tau degradation in primary hippocampal neurons lentivirally transduced with an shRNA against TSG101, an essential component of the ESCRT-I complex. Depletion of TSG101 is a common mechanism for blocking degradation of ESCRT pathway cargo (Edgar, Willen et al., 2015, Maminska, Bartosik et al., 2016), and our shTSG101 hairpin reliably reduces TSG101 levels by ∼60% (Fig. EV2A-B). Using a previously-described cycloheximide (CHX)-chase assay to measure the fold-change in Tau degradation (Sheehan, Zhu et al., 2016), we found that shTSG101 decreased this value by 30% (Fig. 2B-C), indicating its ability to inhibit Tau degradation. Conversely, activation of the ESCRT pathway by TSG101 overexpression increased Tau degradation by ∼30% (Fig. 2D-E). Together, these findings indicate that the ESCRT pathway is both necessary and sufficient for mediating Tau degradation.

**Figure 2.**
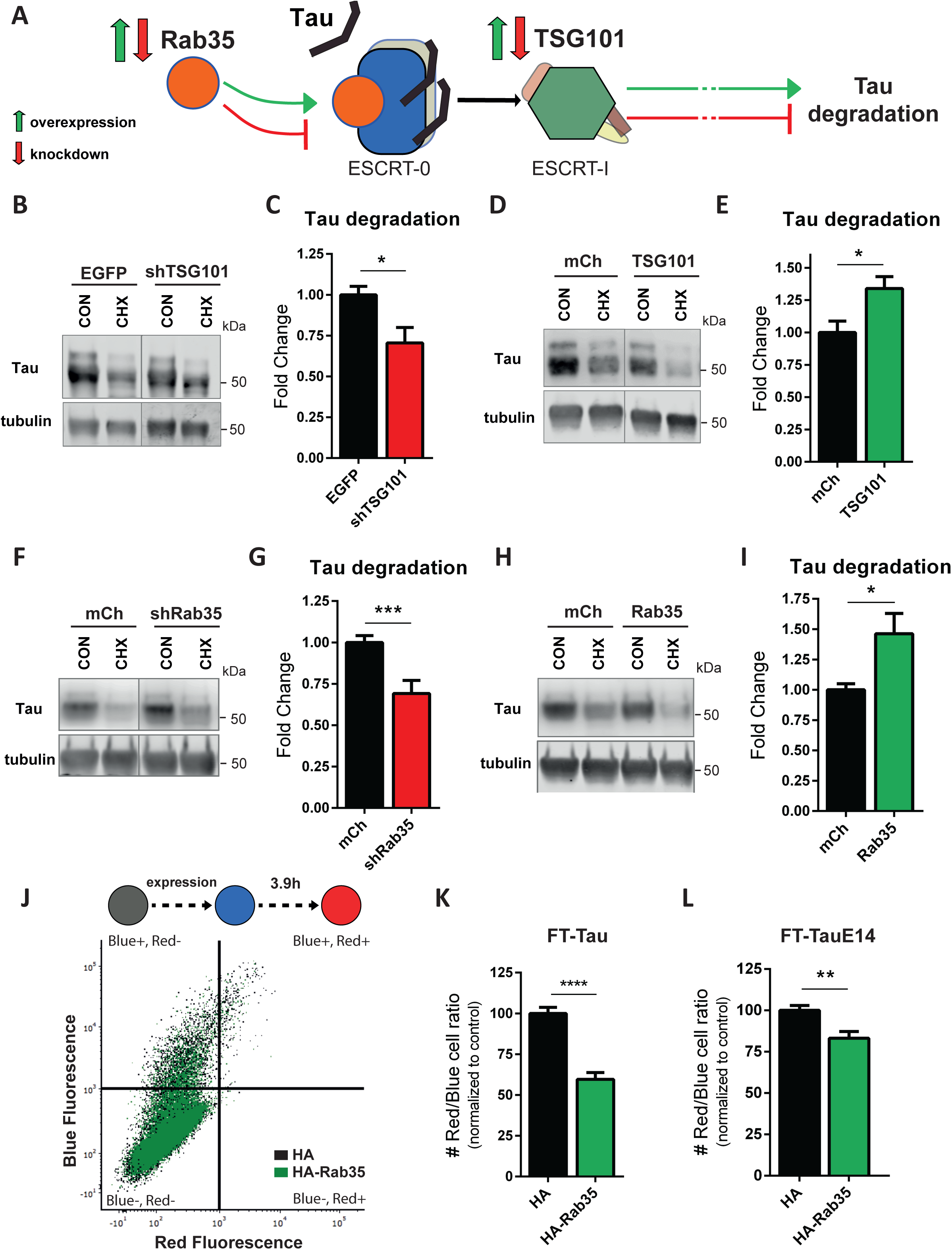
Tau degradation occurs through the Rab35/ESCRT pathway. (A) Schematic diagram of the Rab35/ESCRT pathway, indicating the manipulation of targets (Rab35 and TSG101) used in our studies, and the final readout of Tau degradation; green and red arrows represent overexpression and knockdown of the target proteins, respectively. (B-C) Representative immunoblots (B) and quantification of Tau degradation (C) from 14 DIV primary neurons transduced with EGFP or shTSG101, treated for 24 h with either DMSO (CON) or cycloheximide (CHX), and probed for Tau and tubulin. shTSG101-expressing neurons exhibit markedly decreased Tau degradation compared to EGFP-expressing controls (n=18–19 per condition). (D-E) Representative immunoblots (D) and quantification of Tau degradation (E) from 14 DIV neurons transduced with mCh or mCh-TSG101, treated for 24 h with either DMSO (CON) or cycloheximide (CHX), and probed for Tau and tubulin. Overexpression of TSG101 increases Tau degradation (n=17/condition). (F-GH) Representative immunoblots (F) and quantification of Tau degradation (G) from 14 DIV neurons transduced with mCh or shRab35, treated for 24 h with either DMSO (CON) or cycloheximide (CHX), and probed for Tau and tubulin. shRab35-expressing neurons exhibit markedly decreased Tau degradation compared to mCh-expressing controls (n=23–26 per condition). (H-I) Representative immunoblots (H) and quantification of Tau degradation (I) from 14 DIV neurons transduced with mCh or mCh-Rab35, treated for 24 h with either DMSO (CON) or cycloheximide (CHX), and probed for Tau or tubulin. Rab35 overexpression increases Tau degradation (n=18 per condition). (J) Flow cytometry distribution of N2a cells cotransfected with either HA vector (black) or HA-Rab35 (green) and medium Fluoresence timer-tagged wild-type Tau (FT-Tau); blue fluorescence (y axis) indicates “younger” Tau protein while red fluorescence (x axis) indicates “older” Tau. (K) The ratio of cells expressing red to blue (older:younger) FT-wtTau is reduced in cells overexpressing Rab35, indicating faster Tau turnover (n=9/condition). (L) The ratio of cells expressing red to blue (older:younger) FT-tagged phospho-mimetic E14 Tau (FT-TauE14) is also reduced in cells overexpressing Rab35, indicating that Rab35 also triggers degradation of phospho-mimetic Tau (n=10/condition). All numeric data represent mean ± SEM (unpaired student’s t-test; *p<0.05; **p<0.01; ***p<0.001; ****p<0.0001).

In recent work, we demonstrated that ESCRT-0 protein Hrs is an effector of the small GTPase Rab35, and is recruited by active Rab35 to catalyze downstream ESCRT recruitment and MVB formation (Sheehan et al., 2016). Therefore, we investigated whether Rab35 is also a key regulator of Tau degradation. Again using the CHX-chase assay, we measured Tau degradation in neurons transduced with a previously-characterized shRNA to knockdown Rab35 (shRab35; Fig. EV2C-D)(Sheehan et al., 2016). We found that shRab35 decreased Tau degradation by ∼30% (Fig. 2F-G), indicative of Rab35’s role in this process. Moreover, overexpression/gain-of-function of mCherry-tagged Rab35 increased Tau degradation by nearly 50% compared to mCh control (Fig. 2H-I), demonstrating that Rab35 is a potent regulator of Tau turnover.

We further confirmed the effect of Rab35 on Tau with a flow cytometry assay using N2a cells transfected with HA-Rab35 and medium fluorescent timer (FT)-tagged wild-type Tau (FT-Tau). The emission of medium FT changes from blue to red as it matures, with half-maximal red fluorescence at 3.9 hours (Fig. 2J)(Subach, Subach et al., 2009), and this fluorophore has previously been used to monitor protein degradation based on the intensities of red (“older”) vs. blue (“younger”) fluorescent protein in cells (Fernandes, Uytterhoeven et al., 2014). We found that the ratio of red:blue (old:new) FT-Tau was significantly lower in N2a cells co-expressing HA-Rab35 compared to HA vector control (Fig. 2K), indicating faster Tau degradation. Rab35 overexpression did not alter the red:blue ratio when FT alone was expressed in N2a cells (Fig. EV2E), confirming that Rab35 specifically stimulates Tau degradation. Given that Tau hyperphosphorylation is a key pathogenic event in AD and other tauopathies (Guo et al., 2017), we also evaluated whether Rab35 could stimulate the degradation of hyperphosphorylated Tau, using the phospho-mimetic E14 Tau mutant, in which 14 serine and threonine residues are replaced with glutamic acid (Hoover et al., 2010). The ratio of red:blue FT-E14Tau was again significantly lower in HA-Rab35-expressing N2a cells (Fig. 2L), indicating Rab35’s ability to mediate the degradation of phosphorylated Tau.

Since Tau is phosphorylated at multiple sites, we used antibodies against several phospho-Tau epitopes related to pathogenic Tau activity, including Ser396/404, Ser262, and Ser202, to determine which Tau species were sensitive to Rab35/ESCRT-mediated degradation. Interestingly, we found that degradation of both pSer396/404-Tau and pSer262-Tau (measured by CHX-chase assay) were significantly slowed by knockdown of Rab35, while pSer202-Tau was unaffected (Fig. 3A-B). We saw similar results with knockdown of Tsg101 (Fig. EV2F-G). To further verify that Rab35 stimulates Tau degradation through the ESCRT pathway, we assessed the effect of Rab35 on Hrs/Tau interaction by PLA. Here, N2a cells were co-transfected with FLAG-Hrs and mCherry or mCh-Rab35, and PLA performed with antibodies against FLAG and either total or phospho-Tau epitopes. For total Tau, as well as pSer396/404-Tau and pSer262-Tau, Rab35 expression significantly increased the number of fluorescent puncta per cell (Fig. 3C-D), representing increased interaction of these Tau species with Hrs, and thus their subsequent sorting into the ESCRT pathway. In contrast, Rab35 had no effect on PLA puncta number for pSer202-Tau (Fig. 3C-D). Consistent with our CHX-chase assay findings, these PLA data suggest that Rab35 promotes the degradation of Tau protein through the ESCRT pathway, with some preference for Tau phosphorylated at pSer262 and pSer396/404, but not pSer202.

**Figure 3.**
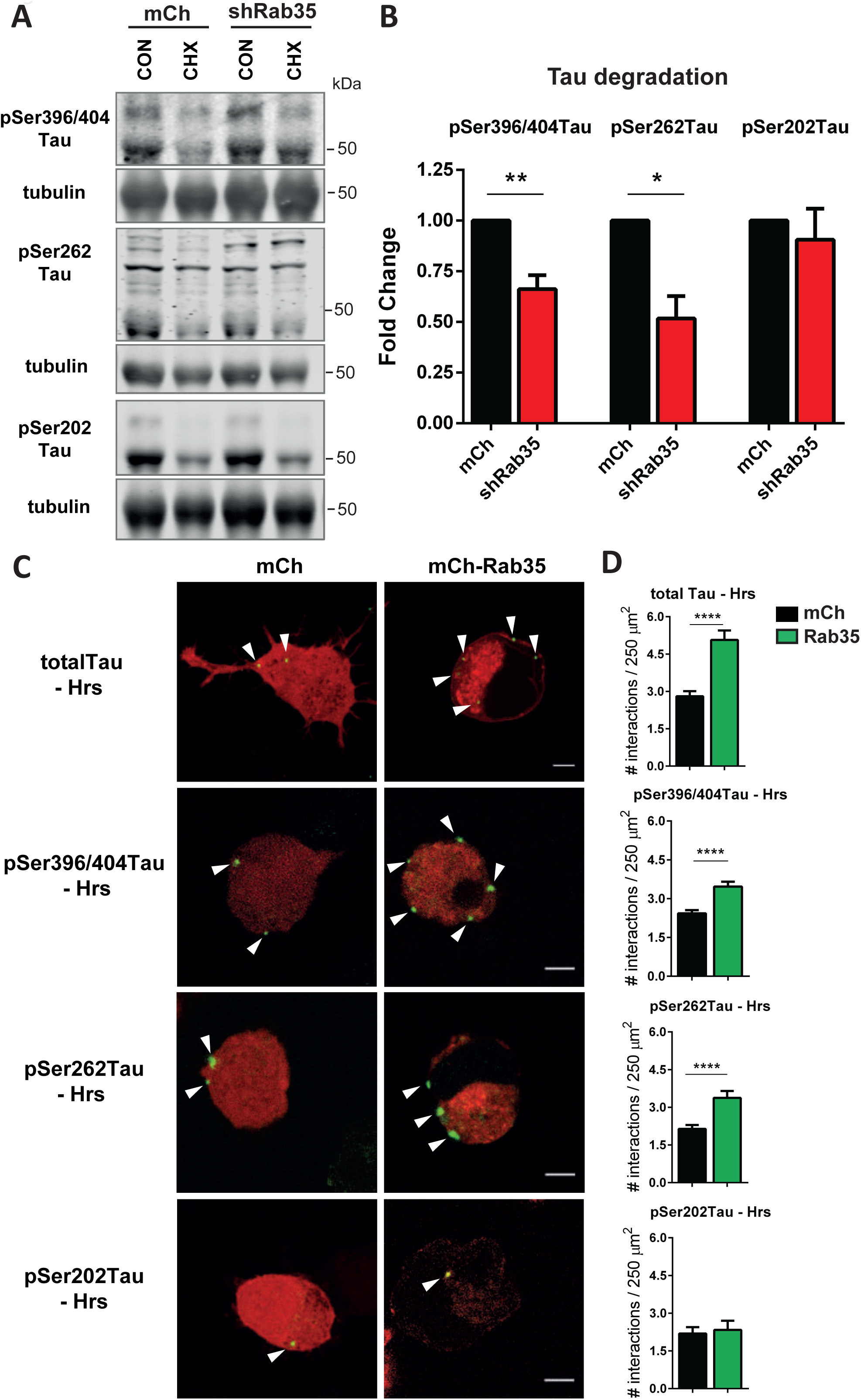
Phospho-dependent selectivity of Tau sorting into the Rab35/ESCRT pathway. (A-B) Representative immunoblots (A) and quantification of Tau degradation (B) from 14 DIV primary neurons transduced with mCh or shRab35, treated for 24 h with either DMSO (CON) or cycloheximide (CHX), and probed for pSer396/404-Tau (PHF1), pSer262-Tau, or pSer202-Tau (CP13) and tubulin. shRab35-expressing neurons exhibit markedly decreased pSer262- and p396/404-Tau degradation compared to mCh-expressing controls, while pSer202-Tau degradation is unaffected (n=4 per condition for pSer396/404-Tau and pSer202-Tau, n=3 for pSer262-Tau). (C) Images of PLA signal (green) for Hrs/Tau interaction in N2a cells co-transfected with FLAG-Hrs and either mCh or mCh-Rab35 (red), and probed with antibodies against FLAG and total (DA9) or phospho-Tau species (pSer396/404-Tau, pSer262-Tau, or pSer202-Tau); scale bar: 5 μm. (D) Rab35 overexpression significantly increases PLA puncta for total, pSer396/404-Tau, pSer262-Tau but not pSer202-Tau (n=121–132 cells for total Tau, 178–195 cells for pSer396/404-Tau, 71–98 cells for pSer262-Tau, 25–29 cells for pSer202-Tau; Mann-Whitney test). Unless otherwise indicated, numeric data represent mean ± SEM (unpaired student’s t-test; *p<0.05; ** p<0.01; ****p<0.0001).

To verify that the Rab35/ESCRT pathway mediates Tau degradation through lysosomes, we next treated hippocampal neurons with bafilomycin (to block acidification and trap lysosome contents) in the presence of mCh-tagged TSG101 or Rab35 to stimulate Tau sorting into this pathway (Fig. 4A). We found that bafilomycin led to a significant accumulation of Tau in lysosomes of neurons expressing either TSG101 or Rab35 versus mCh control, assessed by the fraction of Tau colocalization with LAMP1 (Fig. 4A-B). Furthermore, through sucrose-based fractionation of primary neurons, we found that Rab35 overexpression increased the fraction of Tau in endosomal/lysosomal (Rab35- and LAMP1-enriched) fractions (Fig. EV3), providing further support for the stimulating role of Rab35 on Tau sorting into the endolysosomal pathway. We also evaluated the effects of Rab35 gain-of-function on Tau stability in different neuronal compartments, using CHX treatment combined with immunofluorescence microscopy in hippocampal neurons expressing mCh or mCh-Rab35, as previously described (Sheehan et al., 2016). Although both Rab35 and Tau are enriched in axons, Rab35 overexpression reduced Tau fluorescence in both axons and neuronal somata by between 25 and 50%, demonstrating the ability of Rab35 to stimulate Tau degradation in distinct neuronal compartments (Fig. 4C-D).

**Figure 4:**
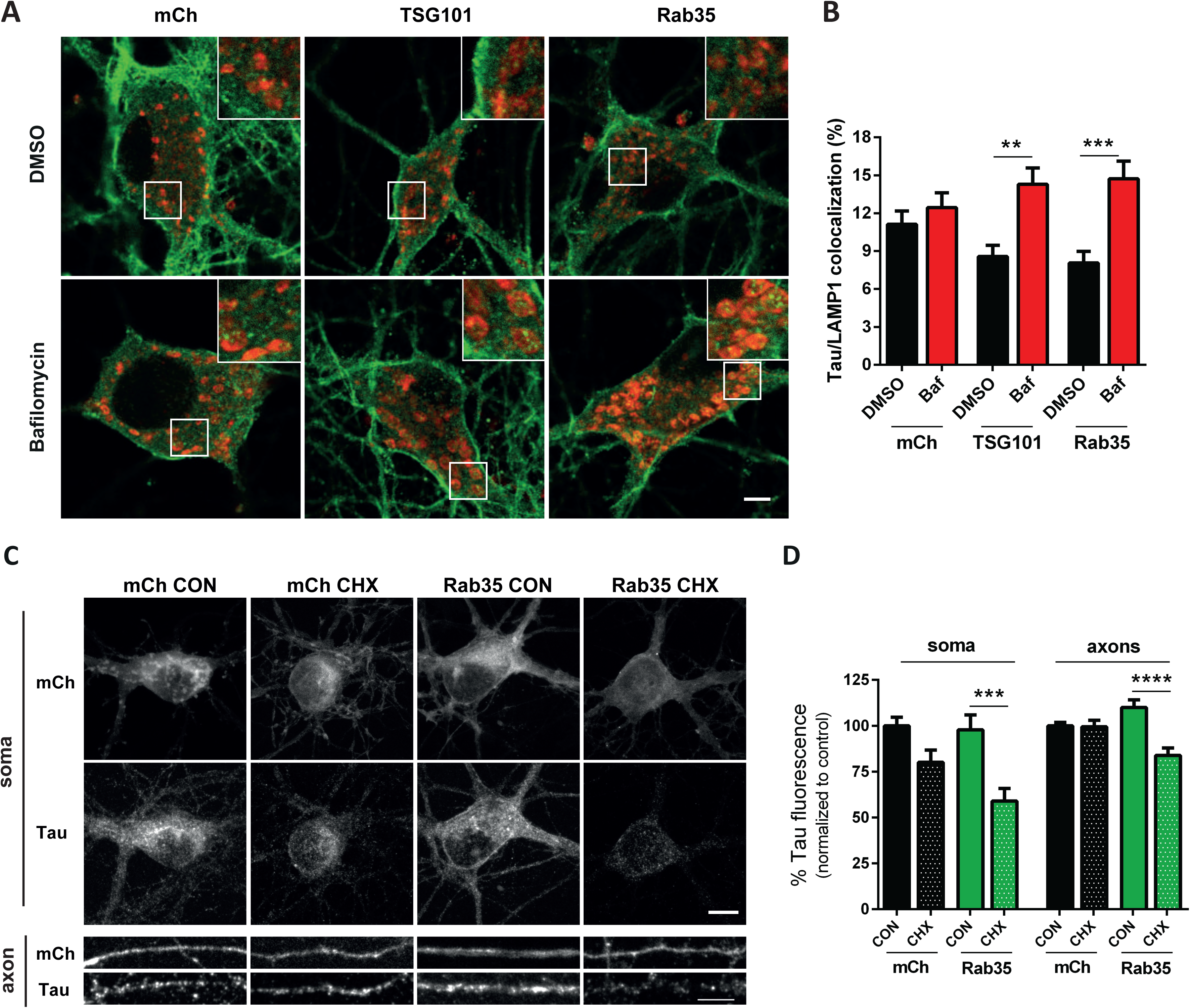
Rab35/ESCRT pathway promotes Tau sorting to lysosomes. (A-B) Representative images and quantification of 14 DIV neurons transduced with mCh, mCh-TSG101, or mCh-Rab35, treated for 5 h with either DMSO (CON) or Bafilomycin (Baf) to block lysosomal degradation, and immunostained for Tau and LAMP1. Overexpression of Rab35 or TSG101 increases Tau accumulation in lysosomes of Baf-treated neurons, indicating that Rab35/ESCRT pathway activation stimulates sorting of Tau into lysosomes (scale bar: 5 μm)(n=25-27 cells/condition; for TSG101, 2-way ANOVA, *Baf* × *TSG101* interaction F_1_, _101_=4,328 p=0.04, overall *Baf* effect F_1_, _101_=10,68 p=0.0015, Sidak posthoc analysis ** p<0.01; for Rab35, 2-way ANOVA, *Baf* × *Ra35* interaction F_1_, _101_=5,36 p=0.02, overall *Baf* effect F_1_, _101_=12,01 p=0.0008, Sidak posthoc analysis *** p<0.001). All numeric data represent mean ± SEM. (C) Images of 14 DIV neurons transduced with mCh or mCh-Rab35, treated for 24 h with either DMSO (CON) or cycloheximide (CHX) and immunostained for Tau; scale bar: 10 μm. (D) Tau fluorescence intensity is markedly reduced in both soma and axons of CHX-treated neurons overexpressing Rab35 compared to mCh, indicating faster Tau degradation (n=24/condition; for axons, n=25-26/condition); for soma, 2-way ANOVA, overall *CHX* effect F_1_, _92_=19.27 p<0.0001, Sidak posthoc analysis *** p<0.001; for axon, 2-way ANOVA, *CHX* × *Rab35* interaction F_1,97_=13.27 p=0.0004, overall *CHX* effect F_1_, _97_=14.34, p=0.0003, Sidak posthoc analysis **** p<0.0001).

### Glucocorticoids decrease Rab35 levels *in vitro* and *in vivo*

Clinical studies suggest that stressful life events and high GC levels are risk factors for AD (Johansson, Guo et al., 2010, Machado, Herrera et al., 2014), and animal studies demonstrate that prolonged exposure to environmental stress and/or elevated GC levels trigger Tau accumulation (Lopes et al., 2016b, Lopes, Vaz-Silva et al., 2016c, Vyas et al., 2016). Based on our findings identifying the Rab35/ESCRT pathway as a critical regulator of Tau degradation, we next examined whether GC treatment altered the levels of Rab35 or ESCRT pathway proteins. Primary neurons were treated with GC and the levels of ESCRT proteins (Hrs, Tsg101, CHMP2b) and Rab35 measured. Interestingly, we found that GC treatment selectively reduced Rab35 protein levels without altering the levels of other ESCRT proteins (Fig. 5A-B). GC are known to regulate gene transcription via activation of the Glucocorticoid Receptor (GR), which binds Gluocorticoid Response Elements (GRE) within the promoter regions of genes (Vyas et al., 2016). We therefore used qPCR to measure Rab35 mRNA levels in hippocampal neurons and N2a cells, and found that GC treatment led to a significant reduction of Rab35 mRNA in both cell types (Fig. 5C). Consistent with the concept that GC regulate Rab35 transcription, we found that the *Rab35* gene contains 14 non-redundant GREs (Table 1). We subsequently measured the levels of Rab35 and other endocytic Rab GTPases in the hippocampi of rats that received GC injections for 15 days (Fig. 5D). The hippocampus displays overt lesions in both stress- and Tau-related pathologies, and is one of the earliest brain regions to show signs of neurodegeneration (Vyas et al., 2016). In line with our *in vitro* findings, Rab35 levels were significantly decreased in GC-injected animals, whereas none of the other Rab GTPases analyzed had significantly altered levels as assessed by immunoblot analysis (Fig. 5E-F). We observed a similar ∼25% decrease in immunofluorescence staining of Rab35 in the dorsal CA1 area of hippocampus in GC-treated versus control animals (Fig. 5G-H; Fig. EV4A). Altogether, these *in vitro* and *in vivo* results suggest that GC specifically decrease Rab35 transcription, leading to reduced Rab35 mRNA and protein levels in hippocampal neurons.

**Figure 5.**
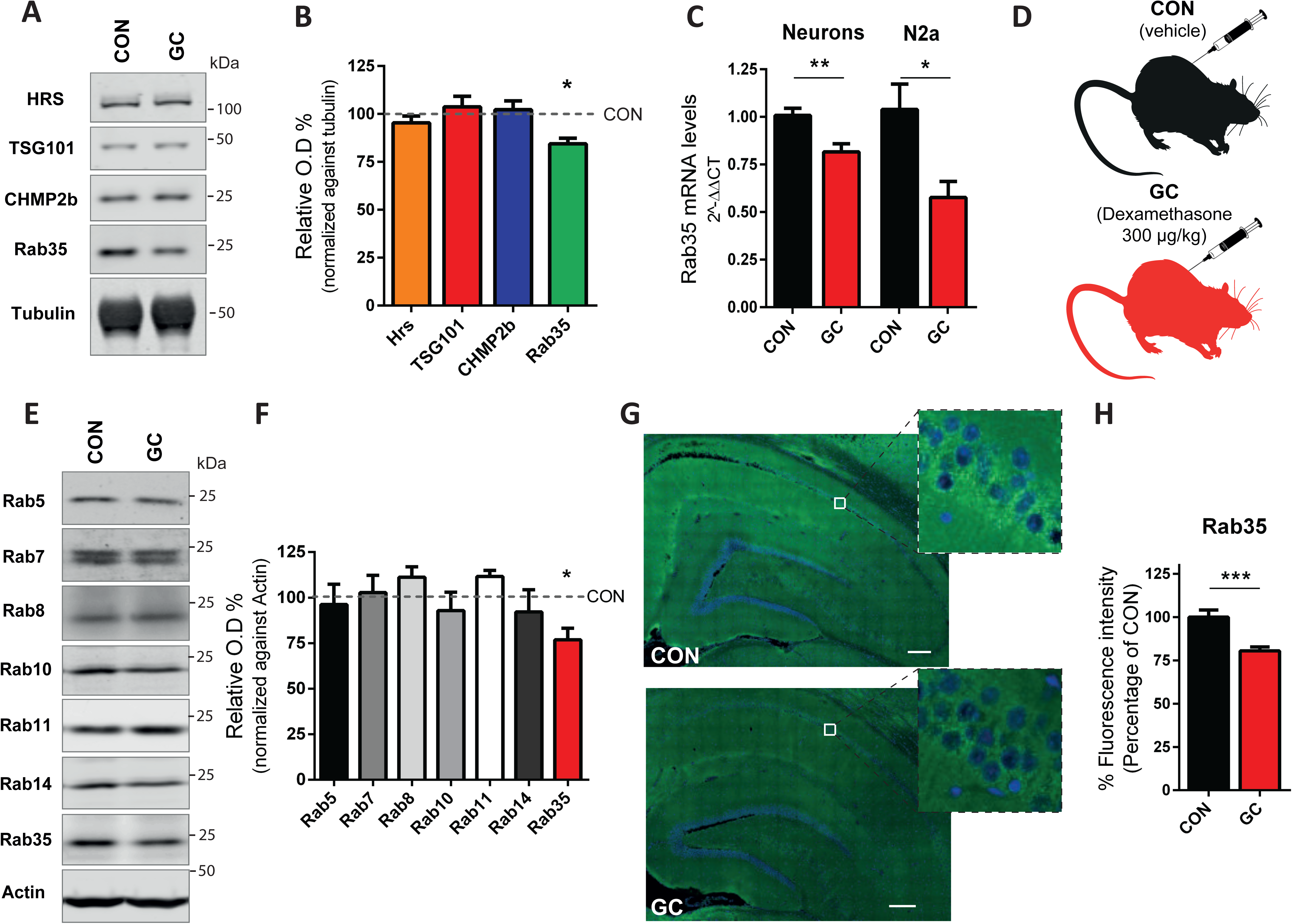
Glucocorticoids decrease Rab35 levels *in vitro* and *in vivo*. (A-B) Representative immunoblots (A) and quantification of Hrs, TSG101, CHMP2b, and Rab35 protein levels (B) from 14 DIV neurons treated with either DMSO (CON) or glucocorticoids (GC). GC treatment selectively decreases Rab35 protein levels without affecting levels of Hrs, TSG101, or CHMP2b (n=9-10/condition). (C) Rab35 mRNA levels are decreased by GC in 14 DIV hippocampal neurons and N2a cells (for neurons, n=9/condition; for N2a cells, n=6/condition). (D) Schematic diagram of GC and vehicle control (CON) treatment in rats for 15 days. (E-F) Representative immunoblots (E) and quantification of levels of different Rab proteins (F) in hippocampus of GC-treated and CON animals. Protein levels of Rab35, but not other Rabs, are decreased in GC-treated animals compared to CON ones (n=5 animals/condition). (G-H) Immunofluorescence staining of Rab35 (green) and DAPI (blue) (G) showing that Rab35 fluorescence intensity is reduced in hippocampal area (CA1) of GC-treated aniamls (H) (n=15 slices/condition; *** p<0.001). All numeric data represent mean ± SEM (unpaired student’s t-test *p<0.05; ** p<0.01; *** p<0.001).

**Table 1.** Chromosomal position of GR binding sites for *Rab35* gene. Our analysis identified 14 distinct glucocorticoid response elements associated with the *Rab35* gene, indicating the relevance of glucocorticoids for regulating *Rab35* transcription.

| **Position** | **Chromosome** | **Start** | **End** |
| --- | --- | --- | --- |
| 1 | chr5 | 115630873 | 115630972 |
| 2 | chr5 | 115631594 | 115631675 |
| 3 | chr5 | 115631971 | 115632003 |
| 4 | chr5 | 115632442 | 115632541 |
| 5 | chr5 | 115632583 | 115632682 |
| 6 | chr5 | 115633507 | 115633606 |
| 7 | chr5 | 115633735 | 115633834 |
| 8 | chr5 | 115635642 | 115635723 |
| 9 | chr5 | 115635734 | 115635827 |
| 10 | chr5 | 115637535 | 115637616 |
| 11 | chr5 | 115640541 | 115640640 |
| 12 | chr5 | 115645909 | 115646008 |
| 13 | chr5 | 115646118 | 115646217 |
| 14 | chr5 | 115647366 | 115647465 |

### Rab35 gain-of-function rescues GC-induced Tau accumulation and neurostructural deficits

Based on our findings that Rab35 mediates Tau turnover, and that exposure to high GC levels downregulates Rab35, we hypothesized that Rab35 overexpression could attenuate GC-induced Tau accumulation and related neuronal atrophy (Green, Billings et al., 2006, Pinheiro et al., 2015, Sotiropoulos, Catania et al., 2011). To test this hypothesis, we performed the CHX-chase assay in hippocampal neurons transduced with either mCh or mCh-Rab35, and treated with GC or vehicle control. As shown in Figure 6, GC treatment reduced Tau degradation in mCherry-expressing neurons, but this effect was completely blocked by Rab35 overexpression (Fig. 6A-B). Similar results were seen in N2a cells, where Rab35 overexpression prevented the GC-driven increase in total Tau levels (Fig. EV4B-C) and in ubiquitylated Tau species (Fig. EV4D-F). Since ubiquitylation is the major signal for cargo sorting into the ESCRT pathway, these findings provide further evidence that Rab35 stimulates the endolysosomal sorting of ubiquitylated Tau.

**Figure 6.**
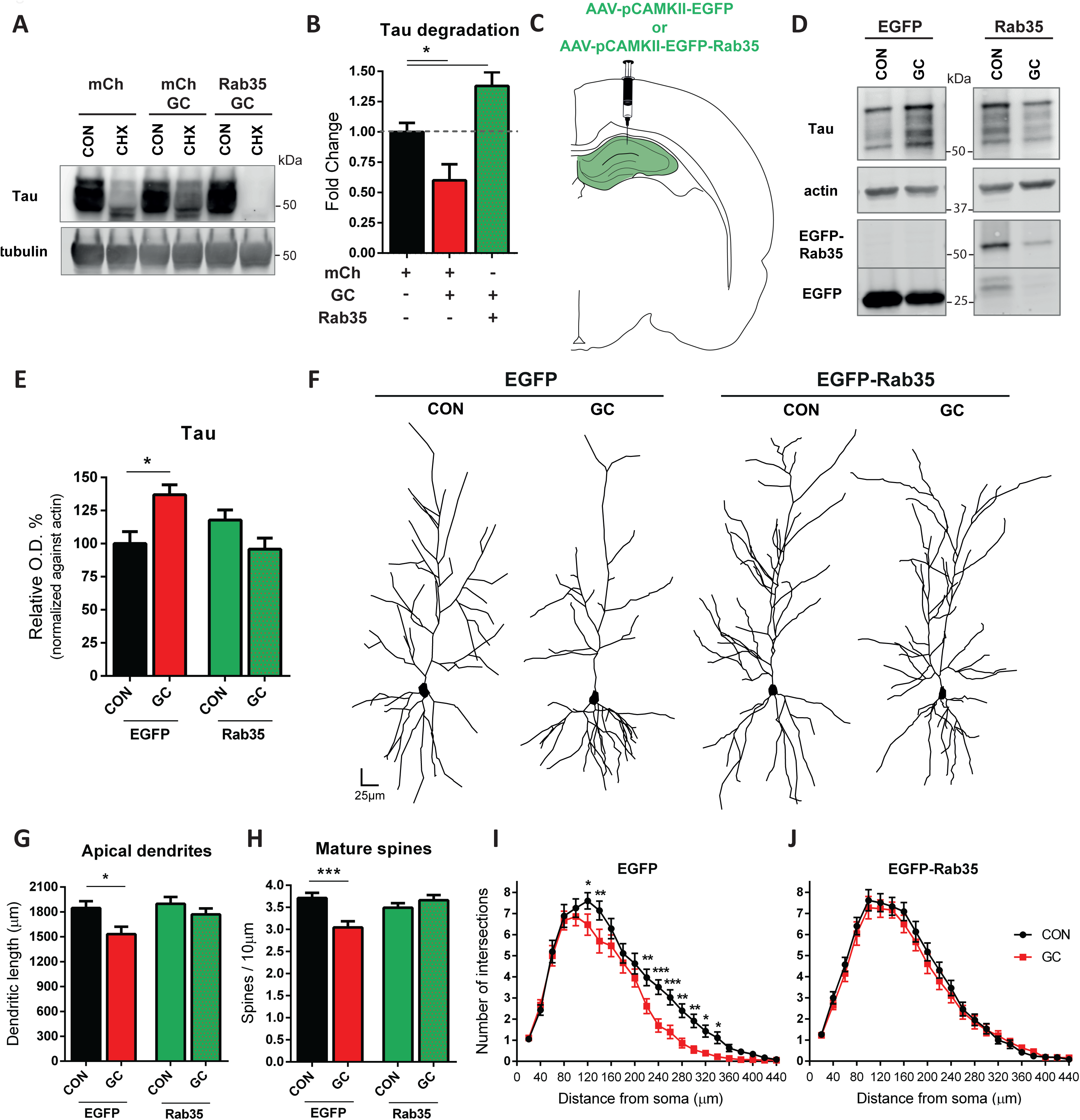
Rab35 overexpression rescues glucocorticoid-induced Tau accumulation and associated neuronal atrophy in rat hippocampus. (A-B) Representative immunoblots (A) and quantification of Tau degradation (B) in 14 DIV neurons expressing mCh-Rab35 or mCh, treated for 24 h with cycloheximide (CHX) or DMSO (CON) under GC conditions. GC significantly decreases Tau degradation, whereas Rab35 overexpression blocks this effect (n=12-14/condition; * p<0.05; 1-way ANOVA, Dunnet post-hoc analysis). (C) Injection of AAV to express EGFP or EGFP-Rab35, driven by the CaMKII promoter, in rat hippocampus prior to GC or vehicle (CON) treatment. (D) Representative immunoblots (D) and quantification of Tau levels (E) in hippocampal synaptosomes reveal that GC increases total Tau levels in animals expressing EGFP, but not Rab35 (n=6 animals/group, 2 replicates, 2-way ANOVA, *GC* × *Rab35* interaction F_1, 39_=10,51 p=0.002; Sidak posthoc analysis *p<0.05). (F) Golgi-based 3D neuronal reconstruction of hippocampal pyramidal neurons (CA1 area). (G) GC treatment reduces the length of apical dendrites in animals expressing EGFP, but not EGFP-Rab35 (n=6-7 animals/group; 6–8 neurons/animal, 2-way ANOVA, *GC* × *Rab35* interaction F_1_, _155_=3.969 p=0.0481, overall *GC* effect F_1_, _155_=8.998 p=0.0031, Sidak posthoc analysis **p<0.01). (H) Mature spine density is reduced by GC treatment in animals expressing EGFP but not EGFP-Rab35 (2-way ANOVA, *GC* × *Rab35* interaction F_1_, _499_=12.33 p=0.0005, overall *GC* effect F_1_, _499_=4.373 p=0.0370, Sidak posthoc analysis * p<0.05; n=6-7 animals/group; 6–8 neurons per animal). (I-J) Sholl analysis of apical dendrites in rat hippocampus shows reduced dendritic intersections after GC treatment in EGFP-expressing animals; however, this effect is not seen in EGFP-Rab35-expressing animals (3-way ANOVA, *GC* × *Rab35* interaction F_1_, _4212_= 14,926 p<0.0001, Simple effect analysis, Sidak test for multiple comparisons *p<0.05 **p<0.01 ***p<0.001, n=6-7 animals/group, 6–8 neurons per animal). All numeric data represent mean ± SEM.

Previous studies showed that exposure to stress or high GC levels induced Tau accumulation, dendritic atrophy, and synapse loss in animals (Lopes et al., 2016b, Pinheiro et al., 2015), and that these hippocampal deficits were Tau-dependent (Lopes et al., 2016a, Lopes et al., 2016b). To test whether Rab35 could protect against these GC-induced effects, we injected middle-aged rats with adeno-associated virus (AAV) to express EGFP or EGFP-Rab35 in excitatory neurons of the hippocampus under control of the CaMKII promoter (Fig. 6C). Animals were subsequently treated with GC or vehicle control. Both EGFP and EGFP-Rab35 injected animals displayed similar body mass loss following GC administration, demonstrating a similar response to high GC levels in the presence or absence of overexpressed Rab35 (Fig. EV5A-C). However, in contrast to EGFP animals that exhibited significant GC-induced Tau accumulation in hippocampal synaptosomes, animals expressing EGFP-Rab35 did not show any such accumulation (Fig. 6D-E). Furthermore, using Golgi-based 3D neuronal reconstruction of CA1 pyramidal neurons in hippocampus, we found that GC significantly decreased the length of apical dendrites in the EGFP control group, but not in animals expressing EGFP-Rab35 (Fig. 6F-G). Notably, EGFP-Rab35 expression alone did not alter apical dendritic length (Fig. 6G), and no difference in basal dendrite length was found between the groups (Fig. EV5D), consistent with previous work showing selective vulnerability of apical dendrites to GC (Lopes et al., 2016b). We also found that GC treatment led to a significant loss of mature spines and concomitant increase of immature spines in EGFP-expressing animals, but no change in mature or immature spine density in EGFP-Rab35 animals (Fig. 6H; Fig. EV5E). GC-induced neuronal atrophy was further confirmed by Sholl analysis, which measures the number of dendritic intersections as a function of their distance from the soma. As shown in Figure 6I-J, GC reduced the number of distal dendritic intersections in neurons expressing EGFP but not EGFP-Rab35. Altogether, these *in vivo* findings indicate that Rab35 overexpression prevents GC-driven neurostructural deficits, implicating Rab35 as an essential regulator of GC-induced neuronal dysfunction.

## Discussion

Impairment of Tau proteostasis is linked to neuronal and synaptic dysfunction in AD animal models and patients (Guo et al., 2017, Ittner et al., 2010, Roberson et al., 2007). Given that Tau accumulation appears to drive neurodegenerative processes in AD and other neurological diseases (see also Introduction), there is growing interest in understanding the mechanisms that mediate Tau clearance, and their selectivity for different forms of Tau. Previous studies have shown that Tau degradation can occur through the ubiquitin proteasome system (UPS), but that macroautophagy plays an important role in the catabolism of aggregated/insoluble Tau, which is not accessible to the UPS (for review, see (Chesser, Pritchard et al., 2013)). Recent work indicates that chaperone-mediated autophagy and endosomal microautophagy also contribute differentially to the degradation of wild-type versus pathogenic mutants forms of Tau (Caballero, Wang et al., 2017), suggesting that Tau turnover is a complex process regulated by multiple factors and involving distinct degradative pathways.

The current study utilizes *in vitro* and *in vivo* approaches to demonstrate a critical role for the endocytic pathway, and in particular Rab35 and the ESCRT machinery, in the turnover of total Tau and specific phospho-Tau species (see Fig. 7). The ESCRT system mediates the degradation of membrane-associated proteins such as epidermal growth factor receptor (Raiborg & Stenmark, 2009), but it has also been implicated in the degradation of cytosolic proteins GAPDH and aldolase (Sahu, Kaushik et al., 2011). These findings are of particular relevance for Tau, which has both cytosolic and membrane-associated pools (Georgieva et al., 2014, Pooler & Hanger, 2010), and has been shown to localize to different neuronal subcompartments based on its phosphorylation state (Hoover et al., 2010, Pinheiro et al., 2015). The current study reveals that Tau appears in Hrs-, EEA1-, and Rab5-positive early endosomes (on the membrane and in the lumen), intraluminal vesicles of MVBs, and LAMP1-positive vesicles and membrane fractions, demonstrating its trafficking through the entire endolysosomal pathway. Moreover, we find that Tau interacts with the initial ESCRT protein Hrs, and that this interaction is strengthened by deubiquitylating enzyme inhibitors, indicating its dependence on Tau ubiquitylation. We also observe that the small GTPase Rab35 is a positive regulator of Tau sorting into the ESCRT pathway. Not only does overexpression of Rab35 stimulate Tau’s interaction with Hrs and subsequent lysosomal degradation, but Rab35 knockdown significantly slows Tau degradation. Interestingly, while Rab35 stimulates the turnover of phospho-mimetic E14 Tau, not all phosphorylated Tau species are equally susceptible to degradation in the Rab35/ESCRT pathway. In particular, we find that pSer396/404 and pSer262, but not pSer202, phospho-Tau species undergo Rab35-mediated degradation, indicative of preferential sorting of specific phospho-Tau proteins into the Rab35/ESCRT pathway. Such differences might reflect changes in ubiquitylation and/or endosomal membrane association of the various phospho-Tau species. Additional work is needed to clarify the relationship between Tau posttranslational modifications *(e.g*. phosphorylation and ubiquitylation) and the sorting and clearance of cytosolic vs. membrane-associated pools of Tau, as well as the impact of stress/GC on these processes.

**Figure 7.**
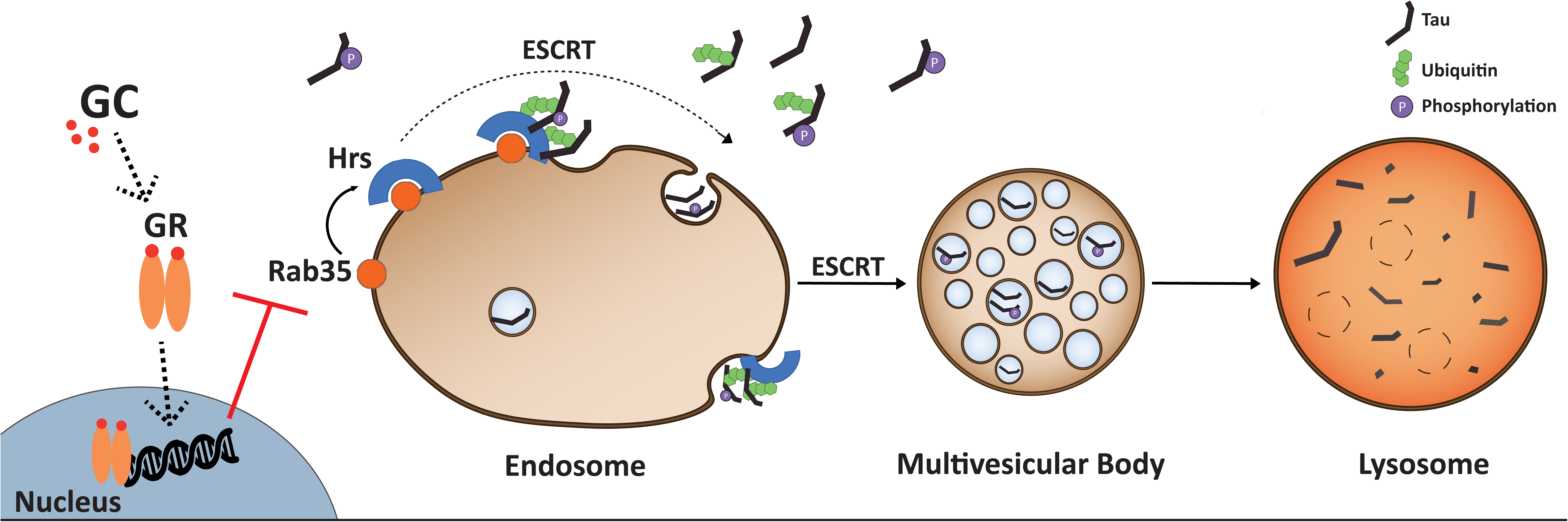
High glucocorticoid levels suppress Rab/ESCRT-dependent Tau degradation, leading to Tau accumulation. (A) Working model of Tau degradation through the Rab35/ESCRT pathway, and its inhibition by glucocorticoids (GC). Rab35 mediates Tau clearance via the endo-lysosomal pathway by recruiting initial ESCRT component Hrs, which recognizes and sorts ubiquitylated Tau into early endosomes for packaging into MVBs. This pathway is impaired by high GC levels, as GC suppress transcription of Rab35, which in turn decreases Tau sorting into MVBs and its subsequent degradation by lysosomes, leading to Tau accumulation and related neuronal atrophy.

We recently showed that Rab35 and ESCRT proteins (e.g. Hrs, CHMP2b) localize to axons and presynaptic boutons, similar to Tau, and that Rab35 stimulates the degradation of synaptic vesicle proteins by mediating the recruitment of Hrs to SV pools in response to neuronal activity (Sheehan et al., 2016). In the current study, we show that Tau interacts with Hrs-positive early endosomes in a Rab35-dependent manner, and that Rab35 stimulates the degradation of both axonal and somatodendritic pools of Tau. Although Tau localizes primarily to axons, many studies (including our own) have shown that it also localizes to the somatodendritic compartment and to dendritic spines under both healthy and pathological conditions, suggesting a synaptic function for Tau (Frandemiche, De Seranno et al., 2014, Ittner et al., 2010, Lopes et al., 2016c, Mondragon-Rodriguez, Trillaud-Doppia et al., 2012, Pinheiro et al., 2015). Moreover, recent work demonstrates more complex intraneuronal trafficking of Tau than previously appreciated, including activity-dependent translocation of Tau to excitatory synapses (Frandemiche et al., 2014) and AMPA/NMDA receptor-dependent Tau hyperphosphorylation and dendritic Tau mRNA translation (Kobayashi, Tanaka et al., 2017). Interestingly, our current findings demonstrate the selective sorting of particular phospho-Tau species into the Rab35/ESCRT pathway, which together with other studies (Frandemiche et al., 2014, Mondragon-Rodriguez et al., 2012, Pinheiro et al., 2015, Pooler & Hanger, 2010) provides novel mechanistic insights into how epitope-specific Tau phosphorylation regulates its trafficking, subcellular localization, and degradation. Our future studies will focus on understanding the role of the Rab35/ESCRT pathway on Tau degradation in different neuronal compartments under pathological conditions, including AD and other tauopathies, wherein Tau is hyperphosphorylated at multiple epitopes (Ittner et al., 2010, Zempel et al., 2010).

Recent evidence suggests that dysfunction of the ESCRT machinery itself is associated with profound cytopathology, as brain-specific deletion of ESCRT components in mice leads to the accumulation of ubiquitylated proteins, impaired endosomal trafficking, and cell death (Oshima, Hasegawa et al., 2016, Watson, Bhattacharyya et al., 2015). Further, a truncating mutation in ESCRT-III protein CHMP2b causes familial frontotemporal dementia and amyotrophic lateral sclerosis, and at the cellular level induces endo-lysosomal pathway dysfunction and the accumulation of ubiquitylated proteins (Clayton, Mizielinska et al., 2015, Zhang, Schmid et al., 2017). Together with our findings, the above studies support the importance of the endo-lysosomal pathway, and specifically the ESCRT machinery, in the clearance of ubiquitylated proteins such as Tau. Indeed, abnormalities of the endo-lysosomal pathway, including endosomal enlargement and high levels of lysosomal hydrolases, are reported as the earliest intracellular features of AD (Nixon & Yang, 2011). Intriguingly, these same features are present in Niemann-Pick Disease Type C (NPC), an inherited lysosomal storage disorder also characterized by Tau pathology (Suzuki, Parker et al., 1995). These shared pathological features of AD and NPC indicate a strong connection between endo-lysosomal dysfunction and Tau accumulation, supporting the importance of this degradative pathway for Tau proteostasis and pathological accumulation.

Multiple cellular pathways are altered by chronic stress, increasing the vulnerability of affected individuals to psychiatric and neurodegenerative diseases such as depression and AD (Ross, Gliebus et al., 2017, Sotiropoulos, Cerqueira et al., 2008). For example, prolonged exposure to stress or high levels of GC trigger Tau accumulation and hyperphosphorylation, accompanied by synaptic missorting of Tau and neuronal atrophy (Green et al., 2006, Lopes et al., 2016c, Pinheiro et al., 2015, Sotiropoulos et al., 2011). Importantly, Tau is essential for this stress/GC-driven damage, as Tau ablation was found to be neuroprotective (Dioli et al., 2017, Lopes et al., 2016c, Pallas-Bazarra et al., 2016). While previous *in vitro* studies showed that GC reduce Tau turnover (Sotiropoulos, Catania et al., 2008), the underlying molecular mechanisms were unclear. The current study demonstrates that GC impair Tau degradation by downregulating Rab35, thereby suppressing Tau sorting into the ESCRT pathway and leading to the accumulation of ubiquitylated Tau (Fig. 7). These results support the concept that ubiquitylation, the major signal for cargo sorting into the ESCRT pathway, represents the first line of cellular defense against Tau accumulation and related neuronal malfunction (Chesser et al., 2013, Kontaxi, Piccardo et al., 2017). Importantly, we find that Rab35 overexpression blocks Tau accumulation and neuronal atrophy induced by high GC levels. Future studies will clarify the potential interplay between endolysosomal machinery and other degradative pathways such as autophagy under stressful/high GC conditions. Our current findings identify the Rab35/ESCRT pathway as a critical regulator of Tau proteostasis, supporting its involvement in the intraneuronal events through which the primary stress hormones, GC, impair neuronal morphology and plasticity in the hippocampus (Sousa & Almeida, 2012). Based on the emerging significance of endosomal trafficking defects in AD brain pathology (Small, 2017), Rab35 and the endolysosomal pathway deserve further investigation for their therapeutic relevance against Tau-dependent neuronal malfunction and pathology.

## Acknowledgments

We thank Dr. Peter Davies (Albert Einstein College of Medicine, NY, USA) for the generous gift of Tau antibodies, Dr. Karen Ashe for pRK5-EGFP-Tau and pRK5-EGFP-Tau E14 (Addgene plasmids #46904 and #46907), and Dr. Ed Boyden for pAAV-CAMKIIa-EGFP vector (Addgene plasmid #50469). We also thank Fidel at the Bronx FedEx station for his help in recovering a valuable package. This work was supported by NIH grants R01NS080967 and R21MH104803 to C.L.W., Portuguese Foundation for Science & Technology (FCT) PhD fellowships to J. Vaz-Silva and T. Meira (PD/BD/105938/2014; PD/BD/113700/2015, respectively), and the following grants to I.S.: FCT Investigator grant IF/01799/2013, the Portuguese North Regional Operational Program (ON.2) under the National Strategic Reference Framework (QREN), through the European Regional Development Fund (FEDER), the Project Estratégico co-funded by FCT (PEst-C/SAU/LA0026/2013) and the European Regional Development Fund COMPETE (FCOMP-01-0124-FEDER-037298) as well as the project NORTE-01-0145-FEDER-000013, supported by the Northern Portugal Regional Operational Programme (NORTE 2020), under the Portugal 2020 Partnership Agreement, through the European Regional Development Fund (FEDER).

## Author Contributions

J.V-S., I.S., and C.L.W. designed the research; J.V-S., M.Z., Q.J., V.Z., P.G., S.Q., T.M., J.S., C.D., C. S-C. and N.D. performed experiments and analyzed data; N.S., I.S., and C.L.W. supervised experiments; J.V-S., I.S., and C.L.W wrote the manuscript.

## Methods

### Primary neurons and cell lines

Primary neuronal cultures were prepared from E18 Sprague Dawley rat embryos and maintained for 14 DIV before use, as described previously (Sheehan et al., 2016). Neuro2a (N2a) neuroblastoma cells (ATCC CCL-131) and HEK293T cells (Sigma) were grown in DMEM-GlutaMAX (Invitrogen) with 10% FBS (Atlanta Biological) and Anti-Anti (ThermoFisher) and kept at 37°C in 5% CO2. During dexamethasone treatment in N2a cells, FBS content in the growth media was reduced to 3%.

### Pharmacological treatments

Pharmacological agents were used in the following concentrations and time courses: cycloheximide (Calbiochem, 0.2 μg/μl, 24 h or 0.1 μg/μl, 16h), bicuculline (Sigma, 40 μM, 24 h), 4-aminopyridine (Tocris Bioscience, 50 μM, 24 h), bafilomycin A1 (Millipore, 0.1 μM, 5h), dexamethasone (Ratiopharm, 10 μM, 48h), dexamethasone (InvivoGen, 20 μM, 72h), epoxomicin (Sigma, 0.1 μM) PR-619 (LifeSensors, 50 μM).

### Lentivirus production, transduction and DNA transfection

DNA constructs were described previously (Sheehan et al., 2016), with the exception of pRK5-EGFP-Tau (Addgene plasmid #46904), and pAAV-CAMKIIa-EGFP-Rab35, which was created by subcloning EGFP-Rab35 into the pAAV-CAMKIIa-EGFP vector (Addgene plasmid #50469). Both pAAV-CAMKIIa-EGFP-Rab35 and pAAV-CAMKIIa-EGFP were then packaged into AAV8 serotype by the UNC Gene Therapy Center Vector Core (UNC Chapel Hill). Lentivirus was produced as previously described (Sheehan et al., 2016). Neurons were transduced with 50–150uL of lentiviral supernatant per well (12 well plates) or 10–40 uL per coverslip (24 well plates) either at 3 DIV for shRNA transduction or 10 DIV in gain-of-function experiments. Respective controls were transduced on the same day for all experimental conditions. Neurons were collected for immunoblotting or immunocytochemistry at 14 DIV.

### Immunofluorescence Microscopy

Immunofluorescence staining in neurons and N2a cells was performed as previously described (Sheehan et al., 2016). Briefly, cells were fixed with Lorene’s Fix (60mM PIPES, 25mM HEPES, 10mM EGTA, 2 mM MgCl2, 0.12 M sucrose, 4% formaldehyde) for 15 min, and primary and secondary antibody incubations were performed in blocking buffer (2% glycine, 2% BSA, 0.2% gelatin, 50mM NH4Cl in 1xPBS) overnight at 4°C or for 1h at room temperature, respectively. For staining of brain slices, coronal vibratome sections (40 μM) of paraformaldehyde-fixed brains were placed in heated citrate buffer for 15 minutes. Sections were then permeabilized using 0.5% Triton X100 for 30 minutes, followed by 5 minutes blocking with Ultravision Protein Block (Thermoscientific). Primary and secondary antibody were diluted in 0.5% Triton/0.2% BSA/0.5% FBS in 1x PBS and incubation was performed overnight at 4°C or for 3h at room temperature, respectively. Images were acquired using a Zeiss LSM 800 confocal microscope equipped with Airyscan module, using either a 63x objective (Plan-Apochromat, NA 1.4), for neurons or N2a cell imaging, or a 40x objective (Neofluar, NA 1.4) for imaging of rat brain sections. Primary antibodies are listed in Table S1.

### Proximity Ligation Assay

Proximity ligation assay (PLA) was performed in N2a cells according to manufacturer’s instructions (Duolink, Sigma). Until the PLA probe incubation step, all manipulations were performed as detailed above for the immunocytochemistry procedure. PLA probes were diluted in blocking solution. The primary antibody pairs used were anti-FLAG (Rabbit; Abcam) and DA9 (anti-Total Tau, Mouse, gift from Peter Davies), anti-FLAG (Rabbit, Abcam) and PHF1 (Mouse), anti-FLAG (Rabbit, Abcam) and CP13 (Mouse), anti-FLAG (Mouse, Sigma) and pSer262 Tau (Rabbit, ThermoFisher Scientific), anti-mCherry (Rabbit, Biovision) and DA9 (Mouse, Peter Davies). Additionally, a blocking FLAG peptide (Sigma), used at a 100 μg/ml concentration, was included to evaluate the specificity of the technique. All protocol steps were performed at 37°C in a humidity chamber, except for the washing steps. Coverslips were then mounted using Duolink In situ Mounting Media with DAPI.

### Tau immunogold staining and electron microscopy

For electron microscope analysis, rat hippocampi were fixed at 4°C with 4% PFA, then transferred to 4% PFA/0.8 % gluteraldehyde in 0.1 M of phosphate buffer (PB) for 1 h and afterwards, to 0.1 M PB. Vibratome-cut axial sections of the dorsal hippocampus (300 μM thick) were collected, and CA1 hippocampal area was surgically removed. Tissue was then carefully oriented and embedded in Epon resin and ultrathin sections (500 Å), encompassing the superficial-to-deep axis, were cut onto nickel grids. For Tau-immunogold staining, sections were treated with heated citrate buffer (Thermo Scientific) for 30 min and then by 5% BSA. Grids were incubated overnight with Tau5 primary antibody diluted in 1% BSA in PB, followed by secondary gold antibody (Abcam). Grids were imaged on a JEOL JEM-1400 transmission electron microscope equipped with a Orious Sc1000 digital camera.

### Image analysis

Images were analyzed and processed using the Fiji software. PLA puncta were counted using the Multi-point tool and cell area was measured with Polygon selection tool. Fluorescence intensity was measured after performing a Z-projection using the SUM function. For cell bodies, the corrected total cell fluorescence was calculated, by subtracting the average fluorescence of the background of the whole area to the Integrated fluorescence density. For Axons, a mask was created in the mCherry channel, representing the experimental condition, using the same threshold value, and the average fluorescence intensity of the axon was then measure in the other channel. Average fluorescence intensity per slice was measured in images acquired from immunofluorescence stained brain slices.

### Coimmunoprecipitacion

For coimmunoprecipitation, N2a cells were transfected with Lipofectamine 3000 according to the manufacturer’s protocol (Life Technologies). Cell lysates were collected 48h after transfection, after 3x washes with cold PBS, in Co-IP lysis buffer (0,1% NP-40, 1mM EDTA in 1x PBS) with protease inhibitor (Roche) and phosphatase inhibitor cocktails II and III (Sigma) and clarified by centrifugation at high speed (10 min, 20,000 rcf). Protein concentration was determined using the BCA protein assay kit (ThermoFisher Scientific) and the same amount of protein was used for each condition. Lysates were pre-cleared using magnetic agarose beads (Chromotek) for 1h at 4°C. For GFP pull down, lysates were incubated with GFP-Trap Magnetic agarose beads (Chromotek) for 3h at 4°C. Beads were washed 3 times with co-IP lysis buffer and then eluted using 2x sample buffer (Bio-Rad) and subject to SDS-PAGE immunoblotting as described below.

### Western Blotting

For Western Blotting experiments, neurons were collected using 2× SDS sample buffer (Bio-Rad). N2a cells were first collected in Lysis Buffer as described previously. Samples were subject to SDS-PAGE, transferred to nitrocellulose membranes using wet or semi-dry apparati (Mini Trans-Blot Cell or Trans-Blot Turbo Blotting System, respectively, BioRad), and probed with primary antibody (Table S1) in 5% BSA/PBS + 0.1% Tween 20, followed by DyLight 680 or 800 anti-rabbit, anti-mouse (Thermo Scientific) or by HRP conjugated secondaries (Bio-Rad). Membranes were imaged using an Odyssey Infrared Imager (model 9120, LI-COR Biosciences) and protein intensity was measured using the Image Studio Lite software (LI-COR Biosciences).

### Cycloheximide-chase fold change calculation

In cycloheximide-chase experiments, the amount of protein remaining after 24h of cycloheximide treatment was calculated as a fraction of the amount of protein in the DMSO treated condition, as previously described (Sheehan et al., 2016). Both levels were previously normalized to the tubulin loading control. The fractional degradative amount of each condition was then normalized to the experimental control by dividing the perturbation condition by the control condition.

### Subcellular fractionation with sucrose step gradient

For each gradient, two T75 culture flaks of primary cortical neurons (7,500,000 cells each) were used. All procedures were carried out at 4°C post-collection and performed as described previously (de Araujo, Huber et al., 2008), with slight modifications. Briefly, neurons were detached from the flask with TrypLE Express (Life Technologies), and washed once with ice-cold Neurobasal and twice with ice-cold PBS. Neurons were then subject to hypotonic shock and allowed to swell on ice for 15 minutes. After cell resuspension in isotonic conditions, neurons were gently lysed with 3x freeze-thaw cycles. The lysates were centrifuged (2,000 × g, 10 min, 4°C), and the post-nuclear supernatant (PNS) was brought to 40.6% sucrose and loaded in the bottom of a centrifugation tube, overlaid with 35% sucrose, followed by 25% sucrose and then 8% sucrose solution (in ddH2O, 3 mM imidazole, pH 7.4 with protease and phosphatase inhibitors). After centrifugation in a Beckman centrifuge (3h, 210,000 × g, 4°C), thirteen 100μL fractions were collected and equal volumes were used for Western blotting, as previously described. Relative protein distribution was calculated after determining the optical density of each fraction and further normalizing against the sum of all fractions. Fractions were divided in different groups according to their respective sucrose gradient.

### Real-time RT-PCR

RNA was extracted from either cortical neurons or N2a cells using TRIzol (Thermo Fischer), and purified using the Direct-zol RNA MiniPrep Plus kit (Zymo Research). RT-PCR was performed in a StepOnePlus RealTime PCR instrument, with iTaq™ Universal Probes One-Step Kit (Biorad), using pre-designed TaqMan probes. Amplification conditions were the following: initial denaturing at 95°C for 10 minutes, 40 cycles of denaturarion at 95°C for 15 seconds and extension at 60°C for 1 minutes. Rab35 levels were normalized to either β-actin (actb) or TATA-binding protein (tbp). Results presented are normalized to β-actin.

### Flow cytometry

N2a cells were detached using TrypLE Express (Life Technologies), for 5 min at 37°C. After washing, cells were resuspended in ice-cold Flow Buffer (0,2% FBS, 0.5mM EDTA in PBS) and strained through a 35μM nylon mesh to promote single cell suspensions and kept on ice. Cells were analyzed in a BD Fortessa (BD Biosciences). Unstained cells were used as a control for background fluorescence. Flow Cytometry data was analyzed using FCS Express 6 (DeNovo Software).

### Ubiquitylation assay

N2a cells were transfected with vectors encoding GFP-Tau and HA-Ubiquitin with either mCherry or mCh-Rab35. After 48h, cells were treated with DMSO vehicle or Dexamethasone, and incubated for another 43h. Cells were then incubated with chloroquine, leupeptin and epoxomicin for 5h. Cell lysates were collected, washed 3x with cold PBS in Lysis buffer (50 mm Tris-Base, 150 mm NaCl, 1% Triton X-100, 0.5% deoxycholic acid) with protease inhibitor (Roche) and phosphatase inhibitor cocktails II and III (Sigma), and clarified by centrifugation at high speed (10 min, 20,000 rcf). Subsequent steps were performed as described in the coimmunoprecipitation section.

### Animals and AAV injection

8–10 month old male Wistar Rats (Charles River Laboratories, Spain; N=5 per group) were paired under standard laboratory conditions (8:00 A.M. to 8:00 P.M.; 22 °C) with ad libitum access to food and drink. Animals received daily subcutaneous injections of the synthetic glucocorticoid, dexamethasone (GC) (300 μg/kg; Sigma D1756; dissolved in sesame oil containing 0.01 *%* ethanol; Sigma S3547) for 14 sequential days, while the other half received daily injections of the vehicle solution (sesame oil with 0.01% ethanol). All experimental procedures were approved by the local ethical committee of University of Minho and national authority for animal experimentation; all experiments were in accordance with the guidelines for the care and handling of laboratory animals, as described in the Directive 2010/63/EU. For AAV injection experiment (N=6-8 per group), animals were anaesthetised with 75mg kg–1 ketamine (Imalgene, Merial) plus 0.5mg kg–1 medetomidine (Dorbene, Cymedica). Virus was bilaterally injected into the dorsal and ventral hippocampus (coordinates from bregma, according to Paxinos and Watson50: −3.0 mm anteroposterior (AP), +-1.6 mm mediolateral (ML), and −3.3 mm dorsoventral (DV) and −6.2 mm AP, +-4.5 mm ML, and - 6.0 mm DV, respectively). A total of 2uL was injected at a rate of 200 nL/minute, and the needle was kept in place for 7 minutes before retraction. Rats were removed from the stereotaxic frame, sutured and allowed to recover for three weeks prior to vehicle or dexamethasone treatment.

### Subcellular fractionation

To obtain the synaptossome membrane fraction, a previously described fractionation protocol was used (Lopes et al., 2016c). Briefly, hippocampal tissue was homogenized [10× homogenization buffer (sucrose 9%; 5 mM DTT; 2 mM EDTA; 25 mM Tris, pH 7.4); Complete Protease Inhibitor (Roche), and Phosphatase Inhibitor Mixtures II and III (Sigma)] and centrifuged (1,000× g). The postnuclear supernatant was subsequently centrifuged (12,500 × g) to yield crude synaptosomal and synaptosome-depleted fractions. The latter was ultracentrifuged (176,000 × g) to yield a light membrane/Golgi fraction (P3) and a cytoplasmic fraction (S3). The crude synaptosomal fraction was lysed in a hypo-osmotic solution and then centrifuged (25,000 × g) to obtain the synaptosomal fraction (LP1).

### Neurostructural analysis

As previously described, half of each rat brain (N=6-7 per group) was immersed in Golgi-Cox solution for 7–10 days. After transfer to tissue protectant solution, vibratome-cut coronal brain sections (200 μM thick) were used. After development, fixation and dehydration, slides were used to perform three-dimensional morphometric analysis. Dendritic arborization and spines were analyzed in dorsal hippocampus (CA1 area). All neuronal dendritic trees were reconstructed at ×600 (oil) magnification using a motorized microscope (Axioplan2, Zeiss) and Neurolucida software (MBF Bioscience). For spine analysis, proximal and distal apical dendritic segments (30 μM) were randomly selected and spines were counted and further classified in immature (thin) and complex/mature (mushroom, wide/thick, and ramified) categories as previously described (Pinheiro et al., 2015). 3 different dendritic segments were selected per neuron.

### *Rab35* glucocorticoid receptor (GR) binding

Binding sites were analyzed with the use of the GTRD database (http://gtrd.biouml.org/)(Yevshin, Sharipov et al., 2017), which is the largest aggregation of ChIP-seq experiment raw data for the human and mouse genome from publically available sources (including GEO, SRA, Encode) using a unified alignment and ChIP-seq peak calling. Peaks were merged into clusters and unique metaclusters (approx. 70M to ensure that transcription factor binding sites are not non-redundant. Our analysis focused on the mouse *Rab35* gene since there is no such comprehensive Chip-seq database for other rodents species. In the search for GR binding sites, we permitted a maximum distance of 1000 nucleotides from *Rab35*.

### Statistical Analysis

Graphing and statistics analysis were performed using Prism (GraphPad). Shapiro-Wilk normality test was used to determine if data sets were modeled by a normal distribution. Unpaired, two-tailed t tests, one-way ANOVA or two-way ANOVAs were used with values of p<0.05 being considered as significantly different.

